# Conditional protein splicing of the *Mycobacterium tuberculosis* RecA intein in its native host

**DOI:** 10.1101/2024.04.15.589443

**Authors:** Ryan F. Schneider, Kelly Hallstrom, Christopher DeMott, Kathleen A. McDonough

**Affiliations:** Biomedical Sciences Department, School of Public Health, State University of New York at Albany; Wadsworth Center, New York Department of Health

**Author notes:** for correspondence: Kathleen A. McDonough, Wadsworth Center, New York State Department of Health, 120 New Scotland Avenue, Albany, NY 12208. authors contributed equally (Ryan F. Schneider, Kelly Hallstrom, Christopher DeMott). College of Saint Rose, Albany NY. **Author Contributions:** RFS, KH, CD and KAM designed experiments. RFS, KH and CD performed experiments. RFS, KH, CD and KAM interpreted data, RFS, KH, and KAM wrote manuscript. **Competing Interest Statement:** The authors have no competing interests.

**Keywords:** Gene expression, post-translational gene regulation, SOS response, DNA damage repair, mobile elements, exaptation

## Abstract

The *recA* gene, encoding Recombinase A (RecA) is one of three *Mycobacterium tuberculosis* (Mtb) genes encoding an in-frame <u>int</u>ervening pro<u>tein</u> sequence (<u>intein</u>) that must splice out of precursor host protein to produce functional protein. Ongoing debate about whether inteins function solely as selfish genetic elements or benefit their host cells requires understanding of interplay between inteins and their hosts. We measured environmental effects on native RecA intein splicing within Mtb using a combination of western blots and promoter reporter assays. RecA splicing was stimulated in bacteria exposed to DNA damaging agents or by treatment with copper in hypoxic, but not normoxic, conditions. Spliced RecA was processed by the Mtb proteasome, while free intein was degraded efficiently by other unknown mechanisms. Unspliced precursor protein was not observed within Mtb despite its accumulation during ectopic expression of Mtb *recA* within *E. coli*. Surprisingly, Mtb produced free N-extein in some conditions, and ectopic expression of Mtb N-extein activated LexA in *E. coli.* These results demonstrate that the bacterial environment greatly impacts RecA splicing in Mtb, underscoring the importance of studying intein splicing in native host environments and raising the exciting possibility of intein splicing as a novel regulatory mechanism in Mtb.

**Significance Statement:** Gene regulation and DNA repair are critical to the success of *Mycobacterium tuberculosis*, a major bacterial pathogen. The present study found significant interplay between the Mtb host environment and splicing behavior of an integrative intein element within the Mtb RecA protein, which is involved in DNA repair. These findings challenge the concept of inteins as strictly selfish genetic elements by showing that activity of the RecA intein in Mtb is finely tuned to its host and raising the possibility that intein exaptation provides Mtb with additional ways to selectively modulate RecA function.

## Introduction

*Mycobacterium tuberculosis* (Mtb) responds to host-associated stresses by tightly regulating its physiological and biochemical processes at multiple levels. Host-associated stressors include hypoxia, starvation, metal toxicity, oxidative stress, and therapeutics during treatment ^1^. These stressors can damage DNA, resulting in replication fork stalling and mutagenesis during DNA repair. Mtb’s primary response to paused replication is expression of the SOS DNA repair regulon, regulated by the LexA transcriptional repressor and Recombinase A (RecA), which is essential for activation of the SOS response ^2^. RecA acts as a coprotease for LexA, which self-cleaves, allowing for increased transcription of SOS genes ^2^, including the error-prone repair polymerase gene *dnaE2*, which is believed to generate the bulk of antibiotic resistance in Mtb ^3^. RecA is also the driver of homologous recombination, a mechanism of double-strand break repair ^4^. While *recA* is dispensable for active Mtb and *M. bovis* BCG infection ^5, 6^, *dnaE2* has been implicated in Mtb survival during active infection ^3^. The effects of an intervening protein sequence (intein) on regulation of RecA production and function within Mtb are unknown, despite a wealth of information on regulation of RecA activity at the transcriptional and post-translational levels ^7–9^.

Inteins are mobile genetic elements comprised of intervening stretches of amino acids that are translated within a host protein ^10, 11^. Inteins must be excised from the precursor protein with consequent ligation of N- and C-terminal regions, termed exteins, for the host protein to be fully functional. Inteins are present in all domains of life but are absent from humans. They typically disrupt essential proteins involved in DNA replication and repair ^10^, leading to interest in their potential as drug targets ^12, 13, 14^. In addition to RecA, Mtb has class I inteins in the DnaB helicase which is essential for replication, and SufB, which is required for iron-sulfur cluster assembly ^15, 16^.

Inteins have been characterized as selfish genetic elements that are maintained because their excision risks causing deleterious mutations of the proteins within which they are found ^17, 10^. However, recent *in vitro* work has demonstrated that intein splicing can be sensitive to environmental triggers, termed conditional protein splicing (CPS) ^18, 19, 20^. Splicing of the *S. cerevisiae* VMA intein is sensitive to temperature and pH ^21^ while splicing of the archaeal *P. horikoshii* RadA intein occurs at high temperatures and in response to ssDNA ^20, 22–24^. These results suggest that some inteins have the potential to benefit their host organisms by providing a means of post-translational gene regulation that can function as an environmental switch. However, mechanisms that affect splicing in native host environments are largely undefined because most intein studies to date have used *in vitro* systems and/or non-native splicing constructs ^25, 21, 26^.

We leveraged the non-essentiality of the RecA intein to study splicing within Mtb bacteria subjected to physiologically relevant conditions to better understand the bacterial and environmental factors that affect intein splicing in a native environment. In contrast to its behavior in surrogate hosts such as *E. coli*, the RecA intein spliced rapidly in TB complex bacteria. RecA splicing was also sensitive to physiologically relevant chemical stressors, including ofloxacin and copper. Stability of the inteins and splicing products within this native Mtb intein environment also differed from previous *in vitro* observations, indicating a critical role for Mtb factors in RecA intein biology. Additionally, we report that a truncated form of RecA that may result from aberrant intein cleavage can activate LexA cleavage in a heterologous host, raising the possibility of an alternate path for selective enhancement of RecA-mediated SOS induction. Together, these results demonstrate significant impact of Mtb factors on native RecA intein biology that contribute to conditional protein splicing (CPS) of an intein in its native environment.

## Results

### Mtb RecA precursor was not detected in Mycobacteria

The RecA protein in Mtb carries a single 48-kDa intein that comprises more than half of the 85 kDa RecA precursor ^16^. The mechanism of class I intein splicing requires a series of reactions involving cysteines, serines, or (rarely) threonines at intein-extein junctions ^10^ (Figure 1A). Cysteines at the C1 and C+1 positions, where C1 is the first position of the intein and C+1 starts the C-extein, facilitate the necessary biochemical reactions for intein excision with concurrent ligation of the flanking exteins (Figure 1A). Non-productive splicing may occur when a trans-acting nucleophile attacks the C1 residue, leading to a cleavage event that liberates the N-extein ^11^.

**Figure 1:**
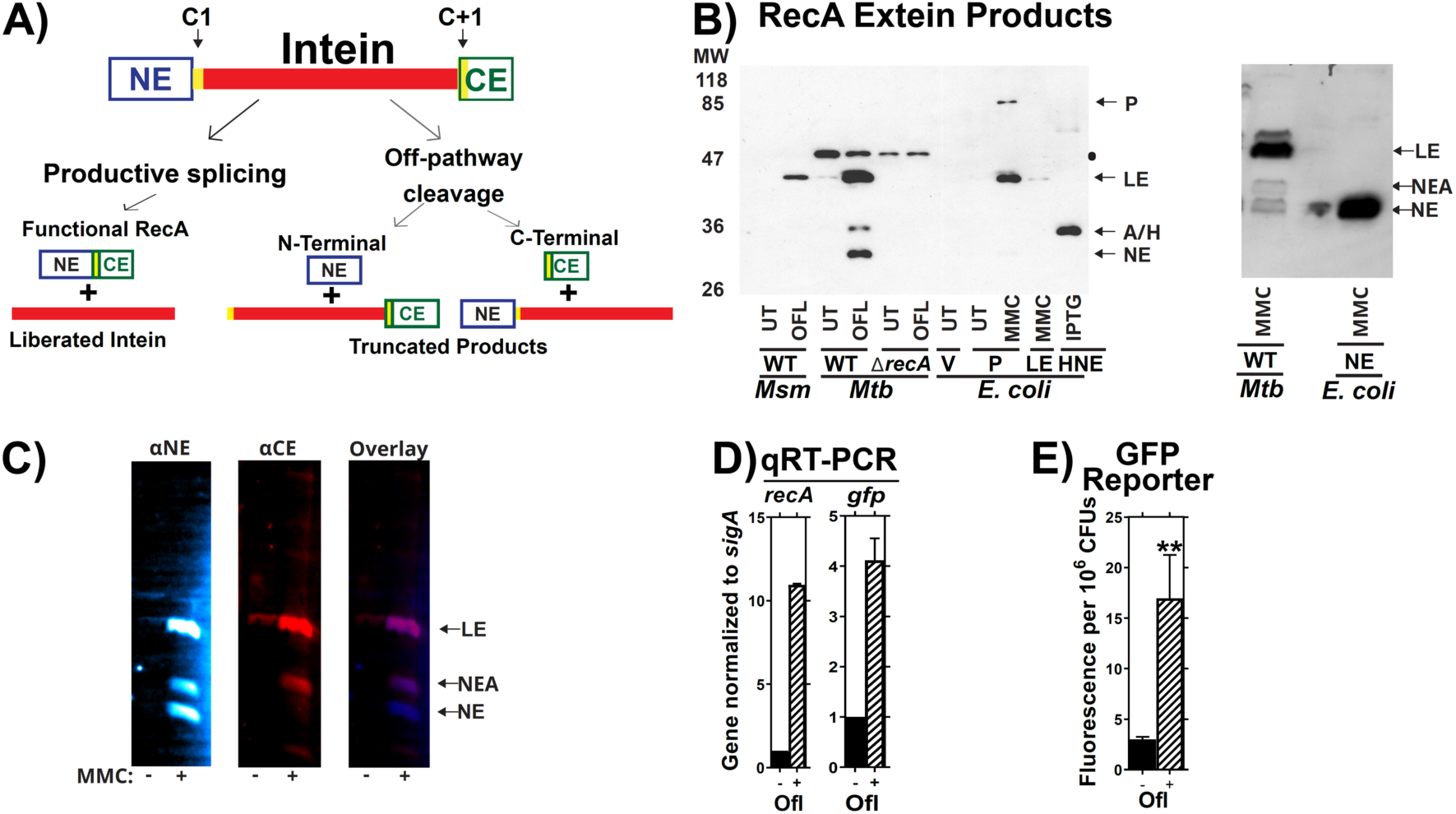
Validation of the splicing reporter system. **(A)** Schematic of RecA splicing products. Mtb RecA is translated as a precursor containing an internal 48 kDa RecA intein. The intein is flanked by an N-terminal extein (NE) of 26.5 kDa and a C-terminal extein (CE) of 11 kDa. Splicing depends on the first residues of the intein (cysteine, C1), and the C-extein (cysteine, C+1). Productive splicing results in precise liberation of the intein from the precursor and allows the N- and C-terminal exteins to be ligated into functional RecA protein (ligated exteins, LE). Alternatively, off-pathway products result in liberation of either the C-extein (C-terminal cleavage) or of the N-extein (N-terminal cleavage). **(B**) Left: Western blot of whole cell lysates from *M. smegmatis* (Msm), Mtb H37Rv mc^2^6230 (Mtb) or recombinant *E. coli* probed with anti- N-extein antibody. RecA splicing was evaluated in response to ofloxacin (OFL), Mitomycin C (MMC), or in untreated cultures (UT) to determine which bands contained splicing or aberrant cleavage products. Msm, which lacks the *recA* intein sequences, was used as a size marker for LE. *E. coli* expressing precursor (P), LE, or NE were included for additional reference bands, as marked, and to compare RecA intein splicing activity in a non-native bacterial host with that in Mtb. RecA was examined in Mtb with a wild-type *recA* locus (WT) or *recA* deletion (Δ*recA*) to identify *recA*-specific bands. A black dot ● denotes a non-specific background band in Mtb cultures. A/H marks the position of a RecA-specific band of unknown identity (called NEA throughout this report) and a 6xHis-tagged N-extein protein (HNE) expressed in *E. coli* BL21 that migrated to similar positions in the gel. Right: *E. coli* expressing a tagless version of the NE was used to further confirm which Mtb band is the NE. Other abbreviations: Vehicle control (V), Isopropyl β-D-1-thiogalactopyranoside treatment (IPTG). **(C)** Lysates of MMC treated Mtb cultures were dual-probed for the (left) NE and (center) CE using fluorescent secondary antibodies. Fluorescent images were overlayed (right) to determine which bands were cross-reactive. The LE and NEA bands were visualized with both antibodies. **(D)** Expression of native *recA* and *recA* promoter driven *gfp* in mc^2^6230 cultures containing the p(*recA*:GFP_v_) reporter was examined via RT-PCR and **(E)** GFP fluorescence in response to ofloxacin. ** indicates p < 0.01 Panel C: n=2. Panel D: n=3.

We measured native RecA splicing in Mtb by performing western blots using antibodies to the N-extein, C-extein, or intein (Figure 1B left). Bands containing specific products of RecA splicing or the excised intein were identified by comparing Mtb lysates with results from Mtb H37RvΔ*recA* and *Mycobacterium smegmatis* (Msm), which has an inteinless RecA used here as a size control for spliced Mtb RecA. *E. coli* expressing the intein-containing full length RecA precursor (P), an inteinless *recA* allele (Ligated Exteins, LE) or a His_6_-tagged RecA N-extein (NE) alone were also used for reference. Bands corresponding to RecA LE were increased in response to the DNA damaging therapeutics ofloxacin (OFL) or mitomycin C (MMC) treatment in Mtb and *E. coli*, consistent with induction of RecA production upon treatment.

Precursor protein was present in an *E. coli* DH5α background (Fig. 1B), but we observed no RecA precursor in Mtb lysates despite testing a range of environmental conditions (not shown). Precursor was also absent when Mtb *recA* was ectopically expressed in Msm (Supplemental Figure 1). This observation is consistent with prior reports describing the absence of intein containing precursor proteins in native environments ^25, 21, 26^ and suggests that the cytoplasmic environment and/or species-specific proteins of the native host bacterium modulate intein splicing.

Additional bands that migrated similarly to NE were also observed. We expressed a tagless NE in *E. coli* and found that it migrated identically to the lower Mtb band identified as NE (Figure 1B right panel). A second NE-associated (NE-A) band was also responsive to MMC treatment in some Mtb cultures that produced RecA. Using a dual probe western targeting both NE and CE, we found that CE antibody was cross-reactive with the NE-A, but not NE, band (Figure1C). We concluded that the lower band corresponds to NE, possibly by off-pathway cleavage, while the upper band is derived from RecA post-splicing. While NE cleavage was observed for the class 3 DnaB intein, DnaBi1, expressed as a reporter construct within Msm ^27–29^, this is the first report of an NE product for a wild-type class 1 intein in its native environment.

### Transcriptional reporter normalizes for recA promoter activity

The absence of precursor made it difficult to quantify relative RecA splicing efficiencies, as previous studies used reporter constructs or purified proteins to measure intein splicing by comparing the ratio of various splicing products to the full-length precursor ^12, 19, 20, 22–24, 29, 30^. We therefore developed a reference system that allows us to measure changes in RecA splicing while controlling for changes in gene expression, including RecA autoregulation. We reasoned that use of a *recA* promoter reporter that expresses the green fluorescent protein venus (*gfpv)* under control of the *recA* promoters on a single-copy integrative vector in Mtb could be used to measure impacts on *recA* transcriptional activity independently of intein splicing.

Our reporter system was validated to ensure that it reflects relative *recA* expression levels in response to DNA damage. qRT-PCR showed that levels of *gfpv* transcripts increased similarly to those of endogenous *recA* after OFL treatment (Figure 1D), confirming that the reporter’s response to environmental stimuli is comparable to that of the native *recA* promoter. GFPv protein levels also increased in response to OFL treatment (Figure 1E), and we concluded that the GFPv reporter serves as a valid readout for changes in *recA* transcriptional activity. RecA protein expression was therefore normalized to GFPv production to account for changes in *recA* transcription that occur during DNA damaging treatments (Figure S2).

### DNA Damage Promotes RecA Splicing

We next used western analyses to measure how RecA splicing is affected by the DNA damaging agents OFL and MMC. MMC is commonly used to induce the SOS response while OFL is a fluoroquinolone, a class of anti-Mtb therapeutics ^25, 31–33^. Intensities of bands containing intein or ligated exteins increased upon treatment with either agent, although levels of excised intein varied more than those of the ligated exteins (Figure 2). In addition to productive splicing that resulted in LE, we observed bands containing NE and the NE-A in response to these treatments (Figure 2B,D). Importantly, these results show that Mtb can produce NE or NE-A in response to specific stress conditions.

**Figure 2:**
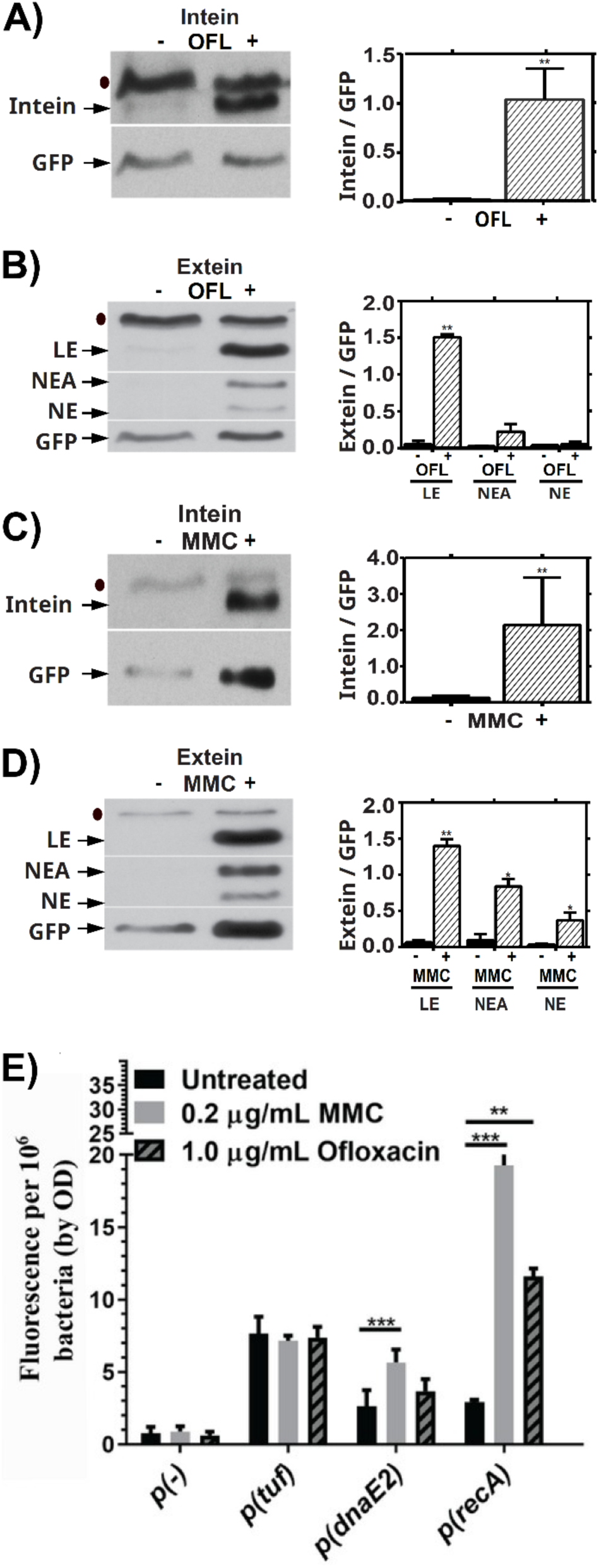
RecA Intein Splicing in Response to DNA Damaging Agents. Presence of native excised intein and ligated exteins in Mtb were measured via western blot following treatment with DNA-damaging drugs ofloxacin **(Panels A, B)** or mitomycin C **(Panels C, D)**. **(A)** Left: Representative western blot of the intein in response to OFL. Right: quantification of 3 biological repeats, normalized to p(*recA*):GFP. **(B)** Left: Representative western blot of LE, NEA and NE in response to OFL. Right: quantification of 3 biological repeats, normalized to p(*recA*):GFP. (**C**) Left: Representative western blot of the intein in response to MMC. Right: quantification of 3 biological repeats, normalized to p(*recA*):GFP. (**D**) Left: Representative western blot of LE, NEA and NE in response to MMC. Right: quantification of 3 biological repeats, normalized to p(*recA*):GFP. **(E)** *recA* transcription in shaking ambient conditions in response to 24 hours of treatment with OFL or MMC as read-out by our GFP based reporter assay. ● indicates background bands in the extein or intein western blot. * indicates p < 0.05, ** p < 0.01, *** p < 0.001, n = 3 biological replicates for A-E.

Transcription of *recA* and the SOS error-prone polymerase *dnaE2* was induced following 24 hours of treatment with either OFL or MMC (Figure 2E), with MMC having a more pronounced effect than OFL on promoter induction. The promoter reporter for a control gene, *tuf*, was unaffected by treatment. Both DNA damaging agents induced similar amounts of ligated RecA relative to our GFPv transcriptional readout (Fig. 2B versus 2D), but the frequency of off-pathway products was most prominent with MMC. This difference suggests that the type of DNA damage may impact production of NE beyond simply induction of RecA splicing. Drugs such as rifampicin and isoniazid that do not damage DNA induced neither *recA* transcription nor RecA splicing after 24 hours of treatment (Supplementary Figure 3). Together, these results suggest that RecA intein splicing is increased by DNA damaging conditions during which more mature RecA protein would be required, as RecA facilitates DNA repair ^34^.

### Oxygen Levels Alter Effects of Copper on RecA Splicing within Mtb

We next asked whether other stress conditions relevant to Mtb pathogenesis also affect RecA splicing. Copper has an antibacterial role in Mtb pathogenesis ^35, 36^ and has been reported to prevent *in vitro* splicing of an Mtb RecA intein reporter protein by directly binding and inactivating the nucleophilic C+1 cysteine ^30^. We tested the effects of copper on RecA splicing within Mtb using our reporter system in ambient air (AMB) or hypoxic (LOC) conditions by growing cultures in these environmental conditions in the presence of 50 µM copper for 7 days. Cultures were grown without oleic acid, albumin, dextrose and catalase (OADC) supplementation as albumin is a known chelator of copper ions ^37^.

We observed no appreciable changes in RecA splicing or transcription in response to copper when Mtb was grown in AMB conditions (transcription not shown). However, RecA intein splicing increased in response to copper in Mtb grown in a LOC environment (Figure 3A, B, LOC lanes, supplemental figure 4). This effect was more pronounced when RecA splicing was measured using ligated exteins than free intein normalized to GFPv (Figure 3B versus 3A). Surprisingly, fully spliced RecA LE predominated when LOC cultures were treated with copper while NE comprised most of the total RecA-associated signal from LOC cultures in the absence of copper (Figure 3B bottom right). This result suggests that copper treatment causes the RecA splicing reaction to shift from NE production to splicing in the native Mtb environment, resulting in more full-length RecA in copper-treated LOC environments. This copper-mediated increase in splicing in Mtb differs from what was observed previously using *in vitro* systems ^30^.

**Figure 3:**
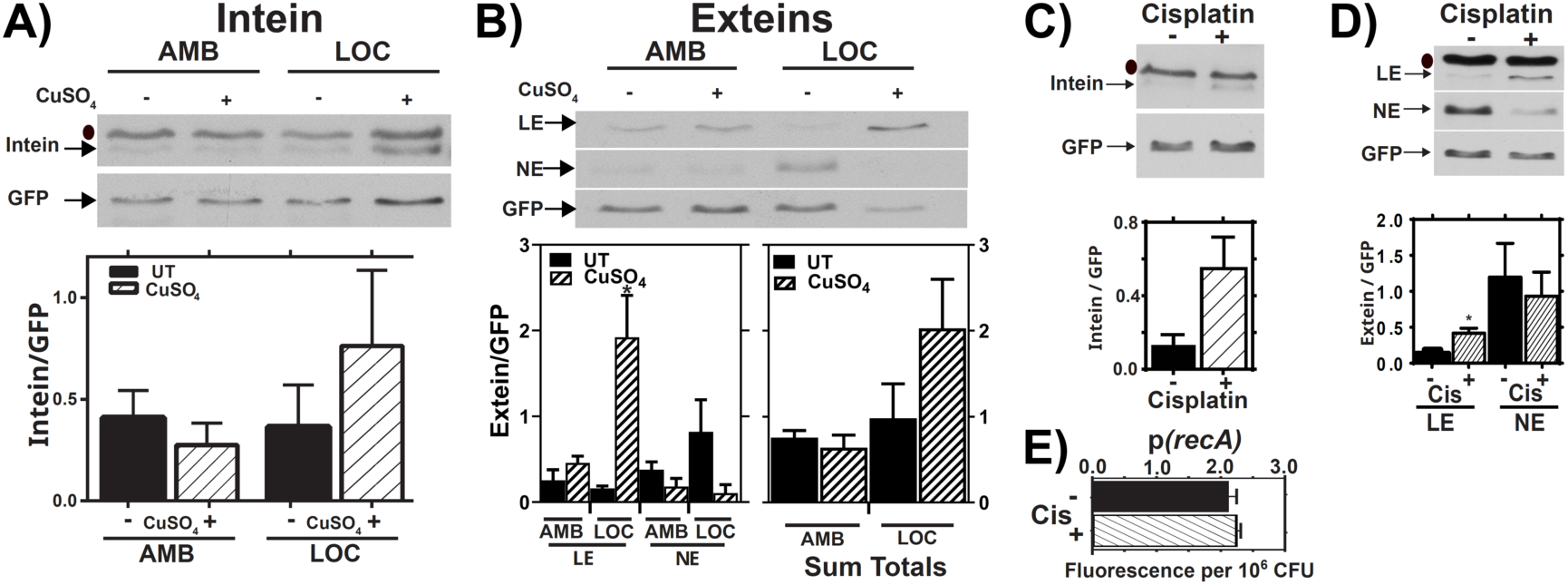
RecA Splicing in Response to Copper and Cisplatin. **(A,B)** Mtb cultures were grown shaking in ambient (AMB) or hypoxic (LOC) conditions in the presence (+) or absence (-) of 50 µM copper sulfate (CuSO_4_) for 7 days. Top: Representative western of intein **(A)** or LE, and NE products in response to copper treatment in AMB or LOC **(B)**. Bars represent mean and standard deviation. Bottom left: Densitometric quantification of LE and NE from 3 biological western blot replicates. Bottom right: Sum of LE and NE bands from the 3 replicates. Bars represent mean and standard error of the mean. **(C,D)** Mtb cultures were treated with cisplatin (CIS), and levels of excised intein **(C)** and LE and NE **(D)** were examined by western blot and normalized to GFP (top). Densitometric quantification of 3 biological western blot replicates. Bars represent mean and standard deviation (bottom). **(E)** *recA* expression in response to cisplatin as read out by our GFP transcriptional reporter assay. ● indicates background bands in the extein or intein western blot, n=3. * indicates p < 0.05.

### Cisplatin Promotes RecA Splicing

The anti-tumor drug cisplatin causes widespread cellular damage, including DNA damage through chromosomal crosslinking, and has been proposed as a potential anti-*Mtb* therapeutic ^38, 39^. A prior report ^12^ showed that cisplatin blocks splicing of a RecA reporter construct *in vitro*, so we tested the effects of cisplatin on native RecA splicing within Mtb. Bacteria were grown to late log phase under AMB conditions and then treated with 3.3 µM cisplatin for 24 hrs. Intein splicing was measured by comparing levels of intein and ligated exteins relative to GFP. We observed a cisplatin-dependent increase in RecA splicing when measured by levels of either the excised intein or RecA LE (Figure 3C, D). The lack of RecA promoter induction in response to cisplatin, as measured by GFP levels from the *recA* promoter-GFP reporter (Figure 3E), suggests that cisplatin toxicity for Mtb also occurs independently of *recA* transcriptional regulation. As with copper, our results suggest that cisplatin can promote, rather than inhibit, RecA splicing in the native Mtb environment, and the basis of cisplatin toxicity is biologically complex.

### The RecA Intein is Not Stable within Mtb

Splicing levels determined by using the intein were often less robust than those determined by using the exteins (Figure 3A). We reasoned that differential stability of these splicing products could cause such an effect, so we examined the stability of native splicing products within Mtb. Cultures were pre-treated with MMC to increase RecA expression and ensure detectable levels of the spliced products. Stabilities of the intein and ligated exteins were measured after halting *de novo* protein synthesis with chloramphenicol (CAM) treatment. RecA intein levels started to decrease by 15 minutes post-treatment and were nearly undetectable by three hours post CAM addition (Figure 4A). In contrast, the level of ligated exteins did not change over the three-hour treatment course (Figure 4B). The instability of the RecA intein may explain why our observations of splicing effects from DNA damage and copper were more pronounced upon measurement of the ligated exteins than the excised intein (Figures 3A,B). These observations indicate that intein stability is regulated by biological host factors within Mtb, a critical observation that was not apparent from prior *in vitro* studies.

**Figure 4:**
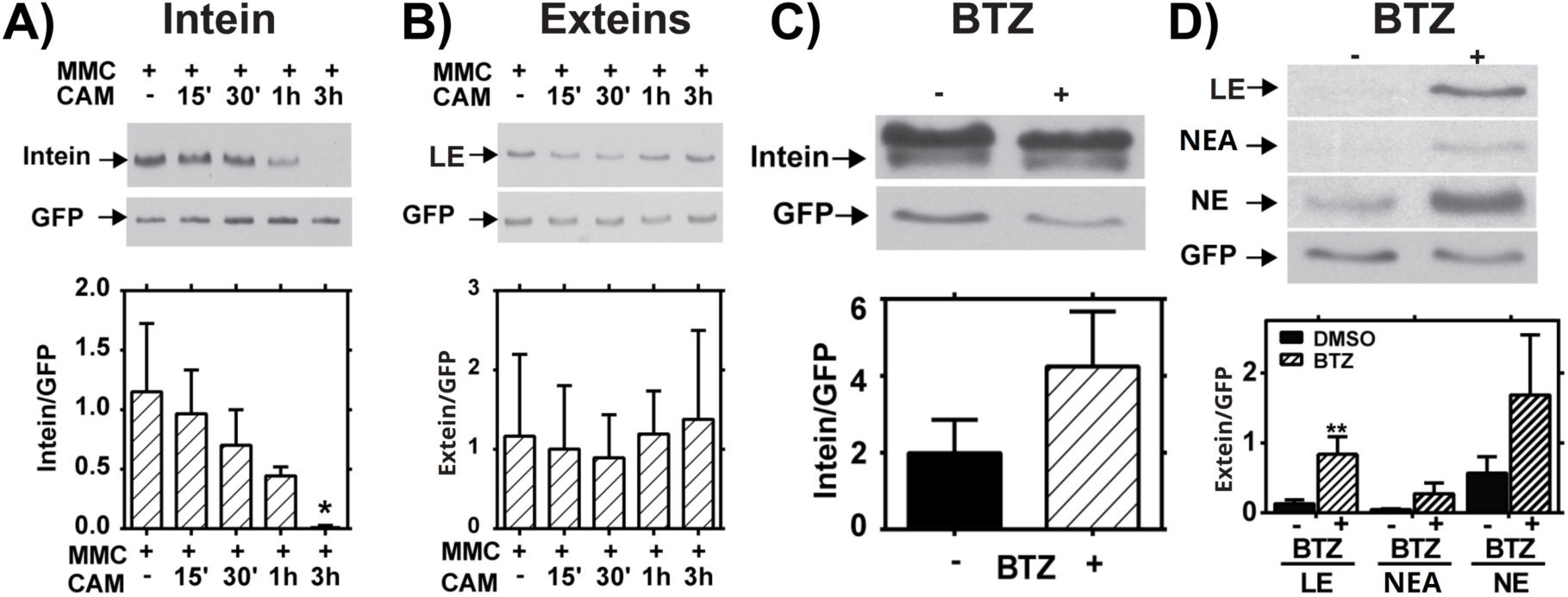
Stability and Degradation of RecA Splicing Products. **(A,B)** Mid-log Mtb cultures were treated with Mitomycin C (MMC) for 24 hours, then treated with chloramphenicol (CAM) for the indicated times to block new protein synthesis. **(A)** Western blot analysis of intein levels over time. Top: Representative blot. Bottom: Quantification of 3 biological repeats normalized to GFP. **(B)** Western blot analysis of ligated extein levels over time. Top: Representative blot. Bottom: Quantification of 3 biological repeats normalized to GFP. **(C,D)** Mtb cultures were treated (+) with bortezomib (BTZ) to block proteasome activity or left untreated (-). **(C)** Western blot analysis of intein levels over time. Top: Representative blot. Bottom: Quantification of 3 biological repeats normalized to GFP. **(D)** Western blot analysis of ligated extein, LE, NEA, and NE levels after BTZ treatment. Top: Representative blot. Bottom: Quantification of 3 biological repeats normalized to GFP; ** indicates p < 0.01.

While prior studies addressed the *in vitro* stability of intein-containing precursors and splicing products ^18, 19^, our results indicated that the stabilities of inteins and ligated exteins are differentially regulated in the Mtb cytoplasmic environment. Mtb encodes a eukaryotic-like proteasome system to which proteins are directed for degradation by the prokaryotic ubiquitin-like protein tag, Pup ^40^. The RecA C-extein contains a putative pupylation site ^41^, so we tested whether RecA stability is regulated by the proteasome. Levels of ligated exteins and free intein were measured following treatment of Mtb with the proteasome inhibitor bortezomib (BTZ) ^42^. BTZ treatment caused accumulation of ligated extein but no significant change in RecA intein levels within Mtb (Figure 4C,D). The free RecA NE and NE-associated bands also accumulated in response to BTZ, although these results were not significant (Figure 4D). Blocking degradation with BTZ did not lead to observation of precursor (not shown). Treatment with protease inhibitors, including those that target serine proteases, had no impact on the levels of LE (Supplemental Figure 5). While degradation by other proteases cannot be ruled out, these data indicate that the proteasome is involved in extein, but not intein, degradation. This result is consistent with recent data assaying RecA stability in proteasome mutants of Msm ^43^. These data further indicate that host factors regulate Mtb RecA intein splicing and stability at multiple levels.

### The N-extein has LexA-activating activity in E. coli

The current paradigm assumes that intein-containing precursors and aberrant cleavage products are non-functional, but recent evidence suggests some precursors can exhibit partial function ^23^. Previous results from Horrii et al ^44^ on *E. coli* RecA led us to compare the functional domains and 3-dimensional architectures of RecA proteins from *E. coli* and Mtb. This analysis suggested that the Mtb RecA N-extein contains the necessary components for LexA activation (Figure 5A, B), which is needed for SOS regulatory function. Potentially functional regions of RecA that are present within the N-extein include the ssDNA binding domains, ATPase domain, and 6 out of 7 monomer:monomer interfaces ^45^. Additionally, the N-extein contains a residue that post-translationally regulates Msm and Mtb RecA function through phosphorylation ^7^.

**Figure 5:**
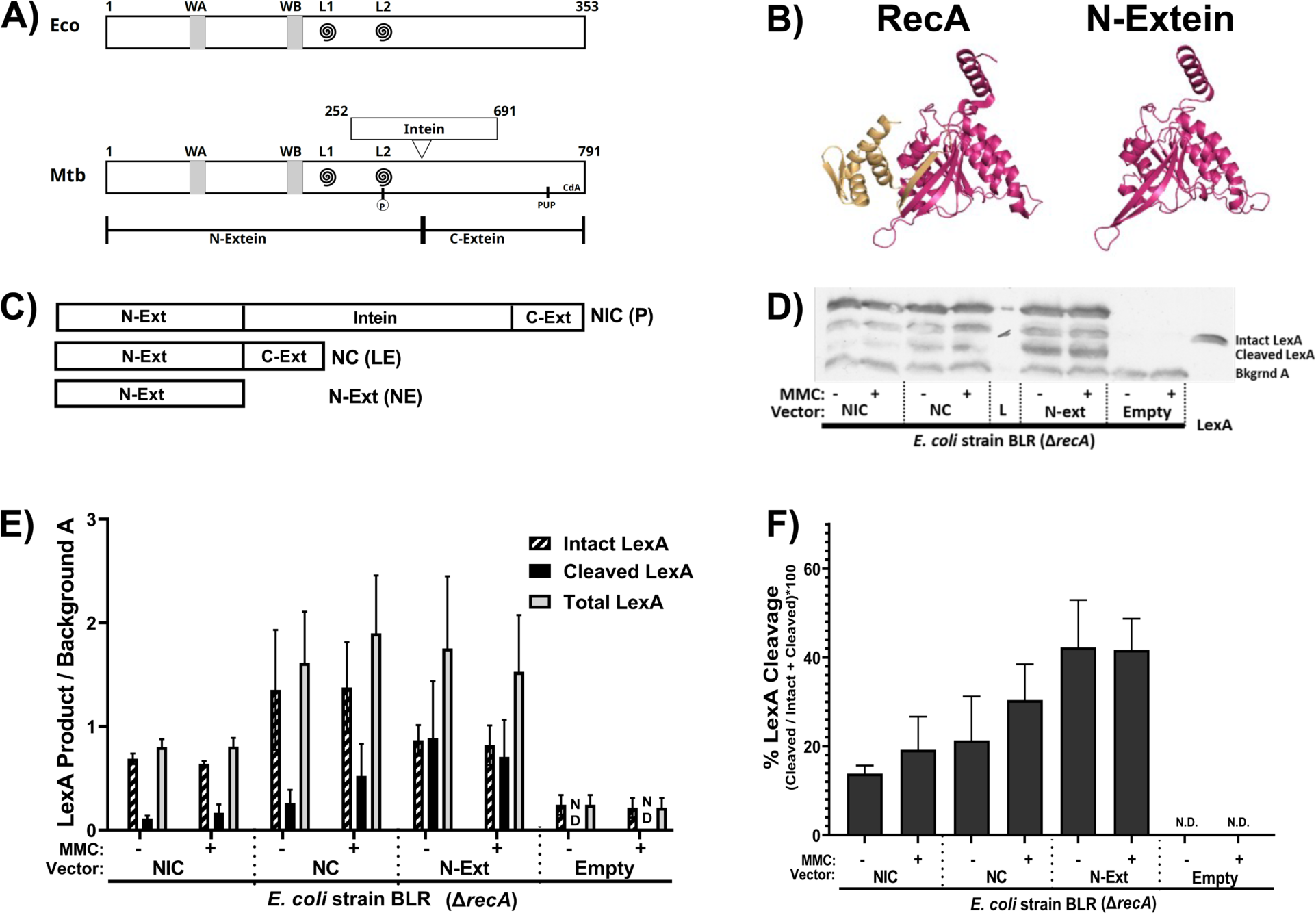
The N-extein product can activate the SOS response in *E. coli*. **(A)** Domain architecture map of (top) *E. coli* RecA protein and (bottom) Mtb RecA protein. WA and WB indicate the Walker A and B motifs. L1 and L2 (swirls) indicate loop 1 and loop 2. The residue in Mtb RecA that can be phosphorylated is in L2. The pupylation site is marked as pup, and the cyclic-di-AMP binding area is labeled CdA. **(B)** (left) Previously crystallized Mtb RecA structure from Datta et al (PDB: 1mo3). The N-extein is colored red, and the C-extein is shaded gold. (right) I-Tasser derived model of folded N-extein colored in red. **(C-F)** Production of LexA and LexA cleavage in E. coli Δ*recA* BLR strains complemented with Mtb RecA constructs driven by the native Mtb promoters: NIC - *recA* including intein; NC - inteinless *recA*; N-Ext - N-extein producing only. **(C)** Schematic of the three constructs. **(D)** Representative western blot probing for LexA. **(E)** Densitometric analysis of LexA western blots normalized to an unchanging background band (Bkgrnd A). **(F)** Quantified cleavage of LexA from western blot analysis. N.D. – Not detected. Error bars indicate SEM of 3 biological replicates.

We tested the ability of the Mtb N-extein to activate the SOS response using *E. coli* strain BLR, a *ΔrecA* derivative of BL21. Three recombinant *E.* coli BLR strains were generated, each complemented with a version of Mtb *recA* driven by the native Mtb *recA* promoter regulatory sequences as follows: 1) NIC complement contains a full-length *recA* gene that includes the intein and generates precursor (P), ligated exteins (LE) and N-extein (NE) in *E. coli*; 2) NC is an inteinless *recA* complement that generates LE; and 3) the NE construct expresses only the RecA N-extein (Figure 5C). LexA production and cleavage were measured via western blot for each of these strains (Figure 5D), as well as in a vectorless control and *E. coli* DH5α (not shown), which expresses a form of RecA that is deficient in homologous recombination but retains the ability to activate the SOS response.

We found that *E. coli* BLR expressing Mtb RecA precursor, inteinless RecA, or the N-extein alone had increased levels of LexA production and cleavage relative to those of the parental *E. coli* BLR *ΔrecA* strain (Figure 5E,F). LexA cleavage was observed in the strain that expressed only the N-extein and occurred in the absence of externally applied DNA damage, demonstrating that the N-extein retains the potential to activate the SOS response.

## Discussion

This study shows that Mtb modulates RecA-intein biology at multiple levels of splicing and product stability. We observed specific effects of the native Mtb host environment on RecA intein splicing and stability that were not apparent from previous studies in non-native and *in vitro* systems, including a lack of precursor accumulation and differential degradation of both intein and extein products within Mtb. Intein splicing within Mtb was stimulated by multiple DNA damaging agents, with production of a semi-functional N-extein protein product in some conditions. Together, these results show that regulation of intein splicing within Mtb is more complex than previously appreciated and raise the exciting possibility that Mtb has adapted to temper the impact of, and possibly even take advantage of, the presence of the RecA intein.

Despite efforts to capture precursor in Mtb, we detected no precursor accumulation in any native Mtb condition tested. Expression of an intein-containing copy of *recA* in *trans* in the normally inteinless Msm also failed to show precursor, despite the clear presence of precursor when *recA* was expressed in *E. coli*. The differences in precursor accumulation in *Mycobacteria* versus *E. coli* suggest that highly conserved *Mycobacteria*-specific factors promote rapid intein splicing or degradation of unspliced precursor. Degradation of ligated RecA by the proteasome was likely due to the presence of a proteasome pupylation site in the C-extein ^41^, and the absence of precursor in BTZ treated samples suggests that rapid degradation alone is unlikely to explain the absence of precursor in mycobacterial cells. Intein degradation may be important for Mtb genome stability, as the intein contains a functional endonuclease ^46–48^ so the mechanism of degradation is of considerable interest. Further study is needed to identify the mycobacterial factors that modulate intein splicing and determine whether the lack of RecA precursor within *Mycobacteria* is due to its instability, rapid splicing and/or a splicing mechanism that occurs co-translationally within Mtb.

The mechanism of N-extein generation likely results from an aberrant cleavage event at the first amino acid of the intein ^11, 49^ (see off-pathway cleavage in Figure 1). However, the present data do not formally rule out proteolytic cleavage at this site, and we are actively working to address this issue. Similarly, the source of NE-A is under investigation. In contrast to N-extein, the NE-A product appears to occur post-splicing, as it includes C as well as N-extein.

The implications for free N-extein function in Mtb are also not known, but it is possible that the N-extein is biologically important. Mtb is a slow-growing microbe with three double strand break repair systems, one of which is preferred over RecA-mediated homologous recombination (HR) when Msm cells are starved to adopt a more Mtb-like growth phenotype ^4^.

N-extein cleavage could separate RecA’s SOS and HR functions. Such separation could allow the SOS response to repair genomic damage during slow-growth or persistence when a second chromosome needed for HR may not be present. Recently, RecA was reported to be essential for resuscitation of persister *Salmonella* cells, presumably because HR repair is required for re-entrance into an actively replicating state ^50^. Retention of the potential to produce full length RecA through intein splicing is consistent with a continuing role for HR in Mtb, as has been shown for *Salmonella* ^50^. DnaE2 has also been implicated in generation antibiotic resistance in Msm persisters ^51^, highlighting the importance of maintaining a functional SOS system during non-replication.

Increased RecA splicing in Mtb exposed to DNA-damaging drugs may provide a post-transcriptional mechanism for generating higher levels of RecA needed in the presence of DNA damage. This possibility is supported by the recent finding that splicing of the archael RadA protein in *P. horikoshii* is stimulated in response to ssDNA, presumably because of DNA damage^23^. Additionally, we found that DNA damaging treatments varied in the extent to which they caused NE accumulation, suggesting that the type of damage that occurs affects levels of both splicing and cleavage. For example, MMC and OFL generated similar amounts of LE relative to transcription, but MMC also increased the amount of aberrant cleavage products (Figure 2). MMC generates crosslinked DNA and reactive oxygen bursts ^52^, while OFL inhibits DNA gyrase, causing areas of ssDNA as well as dsDNA breaks ^53^.

Highlighting the complexity of intein splicing in native biological systems, we found that both copper and cisplatin increased RecA splicing *in vivo*, which contrasts with previous *in vitro* findings ^12^. A prior report showed that CuII blocks splicing of a recombinant RecA splicing reporter protein by directly binding and inactivating the nucleophilic C+1 cysteine ^30^. The basis for the different effects of copper on RecA splicing *in vivo* vs *in vitro* is not clear. CuI is expected to be the predominant species within the intracellular reducing environment of Mtb ^54, 55, 56^, while CuII is the major ion in the previously published *in vitro* system ^30^. Copper can take part in a Fenton-like reaction, generating reactive oxygen species that have potential to damage DNA^57^.

Whether or not the increase in RecA splicing within Mtb occurs as a result of direct interactions between copper and RecA or is due to copper-induced signaling pathways remains to be determined. Therefore, we propose that copper exposure promotes RecA splicing in the native Mtb environment, and the effects of environmental conditions affect different inteins in different ways. We also noted that copper reduced N-extein cleavage as measured by production of NE, but the high level of RecA production in LOC overcame this effect. This oxygen-dependent phenotype may be biologically relevant, as Mtb is likely to encounter copper in microaerophilic conditions during infection.

Our results support the idea that host organisms have evolved to tolerate and possibly exploit intein insertion by using intein splicing as a mode of post-transcriptional gene regulation. In Mtb, for example, this additional layer of gene regulation may allow the bacteria to fine-tune their responses to specific environmental stresses encountered in the host environment, thus supporting population survival. Such adaptation to horizontally acquired genetic material is akin to that observed with pathogenicity islands. For example, *Salmonella* pathogenicity islands (SPIs) are regulated in part by two-component systems located within the core *Salmonella sp.* genome that are sensitive to a range of environmental conditions encountered as the bacteria travel through the gastrointestinal tract ^58, 59^. Similar complex genetic and environmental regulation occurs for virulence genes found on the *Vibrio cholera* pathogenicity island (VPI) ^60, 61, 62^. Thus, it is not uncommon for core genetic elements to evolve into environmentally sensitive regulators of horizontally acquired genetic material in bacteria. It seems likely that the biological and biochemical modes of intein regulation are intricately coupled through direct or indirect mechanisms that are sensitive to environmental conditions. Identification of factors that affect intein splicing in native host environments is critical to understanding evolutionary selection for inteins, the possible benefits and roles of inteins in host organisms, as well as the biology of the organisms themselves.

## Materials and Methods

### Bacterial strains and culture conditions

All strains used are in Table S1. The Mtb mc^2^6230 auxotroph, derived from H37Rv and lacking RD1 and pantothenate synthesis (Δ*RD1,* Δ*panCD);*^63^ ^64^ was grown in Middlebrook 7H9 medium (Difco) supplemented with 0.2% [vol/vol] glycerol, 10% [vol/vol] oleic acid-albumin-dextrose-catalase (OADC), 0.05% [vol/vol] tyloxapol, 0.05% [vol/vol] casamino acids, and 24μg/mL pantothenate. For experiments involving copper, OADC was removed from the medium to avoid chelation of the copper ions by albumin. Fresh cultures were inoculated from frozen stocks for every experiment. Bacteria were typically used after 7 days of growth (late log phase), except where indicated. Cultures were grown at 37°C in 25-cm^2^ tissue culture flasks in auxotroph medium at a shallow depth of 2 mm under ambient (AMB: 20% O_2_, 0% CO_2_) or CO_2_-supplemented microaerophilic conditions (LOC: 1.3% O_2_, 5% CO_2_) in controlled atmospheric incubators, and maintained either shaking or standing.

### DNA cloning and constructs

Plasmids and primers are summarized in Tables S1 and S2. A GFP-based *recA* transcriptional reporter was generated by amplifying the defined *recA* promoter region^8^ with primers (Integrated DNA Technologies) that add BamHI restriction sites each side of the fragment (primers KM4194 and KM4195). The DNA fragment was ligated into pMBC1799, a single copy, integrating vector containing the gene for GFPvenus, that was digested with BamHI. The resulting plasmid, pMBC1809, has the *recA* promoter driving GFPvenus expression. The *recA* promoter consists of the start codon and the adjacent 250bp upstream. This region includes both the P1 and P2 promoters ^8^. pMBC1809 was electroporated into Mtb mc^2^6230.

His-tagged NE or CE-encoding DNA sequences were cloned into the pET28a(+) vector as follows. The full *recA N-extein* was amplified via PCR with EcoRI and HindIII restriction sites included in the primers. The full 771 bp product was cut with EcoRI and HindIII, then inserted into the pet28a+ expression vector. The *recA C-extein* sequence was amplified via PCR with EcoRI and SacI restriction sites included in the primers. The product was cut with the above enzymes, then inserted into the pet28a+ expression vector.

Cloning of *recA* for expression in *E coli* was done as follows. For full length recA, the promoter and open reading frame were amplified from heat killed H37Rv template and cloned into pMBC409, a *Mycobacterial* integrative vector. For inteinless *recA* the promoter + N-extein were amplified together using a reverse primer that added a 3’ overhang homologous to the 5’ end of the C-extein. The C-extein was amplified with a forward primer that added a 5’ overhand homologous to the N-extein. These products were mixed at an equimolar ratio, heated for 5min at 95C and allowed to anneal as the reaction cooled. The resulting product was gel purified and cloned into pMBC409. For N-extein alone, the promoter + N-extein was amplified using a reverse primer that added a stop codon to the end of the N-extein sequence, and this product was subcloned into pMBC409.

### *recA* knockout generation

The *recA* knockout strain was generated in Mtb Mc^2^6230 through phasmid based recombination following Jain et al ^65^. The *recA* phasmid was a gift from Dr. William Jacobs. Briefly, phages were amplified in M. smegmatis containing phasmid. Phages were then used to infect Mtb Mc^2^6230 and bacteria were plated on hygromycin selection media. Individual colonies were picked and expanded in hygromycin containing liquid media. PCR targeting *recA* was done to confirm loss of the *recA* allele. Anti-Reca western blotting was used to further confirm that RecA was not produced in this strain.

### Treatment conditions

#### Mitomycin C, Ofloxacin, Nitric Oxide and Cisplatin

Mtb mc^2^6230 cultures were grown with gentle shaking in AMB or LOC conditions for 6 days to late log phase, then subjected to various treatments for 24 hrs. Nitric oxide treatment was done by using a final concentration of 250µM DETA-NO (Caymen Chemicals). Final drug concentrations were for 0.2μg/mL for mitomycin C (Fisher Bioreagents), 20ng/mL for cisplatin (Sigma-Aldrich), and 1µg/mL for ofloxacin (Sigma-Aldrich). Ofloxacin stock solutions were made at 5mg/mL in 20µM NaOH, so 20µM NaOH lacking ofloxacin was used as a vehicle control for ofloxacin experiments.

#### Copper treatment

Mtb mc^2^6230 cultures were grown in the absence of OADC with or without 50µM CuSO_4_ for 7 days shaking or standing in AMB or LOC conditions. To control for effects of copper on growth, cultures were also grown up to 1 OD in standard auxotroph media, then resuspended in media lacking OADC and treated with 50μM CuSO_4_ for 7 days. Cultures were grown shaking or standing in AMB or LOC conditions. Where indicated, samples for OD_600_ were taken and measured on a Sunrise 96-well plate reader (Tecan).

### Inhibition of proteasome

Mtb mc^2^6230 cultures were grown as previously described ^66^ for 4.5 days to mid-log phase, then treated for 48 hrs with 40μM bortezomib (BTZ, Sigma-Aldrich) dissolved in DMSO^66^. DMSO was used as a vehicle control.

### Chloramphenicol time course

Mtb mc^2^6230 cultures were grown for 3 days at shaking/AMB at 37°C then treated for 24 hrs with 0.2µg/mL of MMC to induce expression of RecA as previously described ^8^. Subsequently, cultures were treated with 35μg/mL CAM (Sigma-Aldrich) to block *de novo* protein synthesis for 15, 30, 60, or 180 min.

### Antibodies against RecA domains

Polyclonal antibody sera from mice raised against the NE were generous gifts from Dr. Guangchun Bai. Anti-intein polyclonal antibody sera raised from rabbits (using Titermax adjuvant) were generous gifts from Dr. Marlene Belfort. Purified C-extein was covalently linked to keyhole limpet hemocyanin using the Imject EDC mcKLH spin kit (ThermoFisher) for antibody production due to its small size (11kD). Polyclonal rabbit antibodies were generated by Pocono Rabbit Farm and Laboratory Inc using Titermax adjuvant. Bleeds were screened for cross-reactivity and sensitivity, and the first final bleed was used for further western blot analyses.

### Mycobacterial protein extraction

Liquid cultures of mc^2^6230 were grown as described to the desired timepoints after treatment. The bacteria were then pelleted and washed with cold PBS containing 0.25% protease inhibitor cocktail (Sigma-Aldrich). The pellet was resuspended in lysis buffer containing 0.3% SDS, 28 mM Tris-HCl, 22mM Tris-Base, and 1% protease inhibitor cocktail. Proteins were extracted by vigorous sonication and freeze/thaw cycles. Cell debris was removed by centrifugation, and the resulting supernatant was used as total protein lysate. Lysate concentrations were determined using a NanoDrop system, and samples were normalized to the same concentration using lysis buffer. Normalized samples were then mixed 1:1 with 2x loading dye containing 40% TCEP and denatured at 95C for 5 min prior to loading onto 12% (ligated RecA exteins) or 8% (RecA intein) denaturing SDS-PAGE gel. The gels were run for 50 min at 100V. Proteins were then blotted onto PVDF membrane using a standard wet-transfer protocol. Equal loading of samples was visualized using a duplicate 12% SDS-PAGE gel stained with standard Coomassie Brilliant Blue staining protocol.

### Western blotting to detect RecA products

To detect RecA or RecA intein, the membrane was blocked with TBST + 5% milk. After blocking, the membrane was probed with mouse-anti-RecA or rabbit-anti-RecA intein polyclonal serum at a 1:3000 dilution in TBST + 0.2% milk, then probed with goat-anti-mouse or goat-anti-rabbit HRP antibody at a 1:5000 dilution. To detect GFP, membranes transferred from the 12% SDS-PAGE gel were stripped then probed with mouse-anti-GFP antibody (Invitrogen), then with goat-anti-mouse HRP antibody (Jackson ImmunoResearch). Peroxidase detection was carried out using the SuperSignal^TM^ West Pico and West Femto Chemiluminescent Substrates (Life Technologies). Densitometry was performed using FIJI (NIH). Where indicated, the ligated extein or excised intein signal was ratioed to that of the corresponding GFP signal, which was used as an internal reference control for changes in *recA* transcription and expression. Students’ T-test was used to evaluate such results from at least three independent biological replicates, in which p<0.05 was considered statistically significant. Error bars represent standard deviation of the mean.

For dual probe western blots, the protocol remained the same except as follows: Blocking was done in TBST + 7.5% goat serum (GIBCO). The primary antibody incubation step included both mouse anti-N-extein and rabbit anti-C-extein antibodies (polyclonal sera) at a 1:3000 dilution each in TBST + 0.3% goat serum. The secondary antibody incubation step included Alexa-488 conjugated goat anti-mouse IgG and Alexa-647 conjugated goat anti-rabbit IgG (Jackson ImmunoResearch) diluted 1:5000 each in TBST + 0.3% goat serum. Membranes were imaged on a BioRad Pharos detection system using the Alexa-488 and Alexa-647 channels.

### RNA extraction

Liquid cultures of mc^2^6230 were grown in the previously described conditions. Flasks were immediately treated with 500μM guanidine isothiocyanate (GTC) upon removal from incubator. The cultures were pelleted, and total bacterial RNA was extracted using TRIzol^®^ reagent (Life Technologies) per manufacturer’s instructions. Briefly, following resuspension in 1mL of TRIzol reagent, cells were lysed mechanically with three rounds of bead beating (BioSpec Products), 70s each, in a mixture of 0.1mm zirconia-silica beads (BioSpec Products). The supernatant was transferred to a fresh microcentrifuge tube and allowed to incubate at room temperature for 5 min to allow for nucleoprotein complex disassociation. 300μL of chloroform were added to the supernatant, then vortexed for 15-20s. The samples were then centrifuged at 13,000 rpm for 15 min at 4C. The upper aqueous phase was removed, and nucleic acid was precipitated with isopropanol at room temperature. RNA was pelleted and washed with ice cold 75% ethanol and resuspended in RNase-free water. DNA was removed with RNase-free DNase (Qiagen) in solution following the manufacturer’s protocols (Qiagen). RNA was assessed for DNA contamination using PCR and RNA integrity was assessed by electrophoresis.

### qPCR

cDNA was prepared from quality RNA for quantitative PCR (qPCR). Briefly, 0.5μg of RNA was used to generate cDNA using random hexamer primers and RT Superscript III following the manufacturer’s protocol (Invitrogen). The qPCR was performed using primers that were validated for efficiency. qRT targets examined were native RecA, and GFPv. qPCR was performed on a 7500 Real Time PCR System (Applied Biosystems) and analyzed using the comparative *C*_T_ method, normalizing transcripts of interest to the 16S-rRNA transcript ^67^. Error bars represent standard deviation of the mean of at least two independent biological replicates.

### LexA cleavage assay

The effects of Mtb RecA N-extein on LexA production and cleavage were assayed in *E. coli* Δ*recA* strain BLR (BLR) expressing derivatives of the Mtb *recA* gene driven by native Mtb *recA* promoter regulatory sequences. Strains included: BLR::vector only control; BLR:: N-extein of *recA* (N); BLR:: precursor *recA*, which includes the inein (NIC) and BLR::inteinless *recA* (NC). Purified *E. coli* LexA protein was purchased from Abcam to use as a western blotting size control. Cultures were grown overnight, and then diluted to 0.5OD_620nm_ and split into duplicate. One of the duplicate cultures was treated for 75minutes with 200ng/mL MMC, and the other was left untreated. Protein was harvested and separated via SDS-PAGE to western blot against LexA (antibody purchased from Active Motif). Intensities of intact and cleaved LexA bands were normalized to a background band that was unaltered by treatment using FIJI.

### Modeling RecA N-extein folding

To predict N-extein folding, we utilized the iTasser server ^68–70^. We used previously solved structures of *E. coli* RecA filamenting on ssDNA, or *M. smegmatis* RecA (monomer) ^71^ as a template.

### Fluorescence plate assay

OD_620nm_ of cultures was first measured in a Sunrise 96-well plate reader (Tecan). Fluorescence of cells carrying the P*recA*-GFPv reporter was then measured in an Infinite F200 96-well plate reader (Tecan) using a filter cube with excitation range of 475-495nm and emission range of 512.5-537.5 nm. Subsequently, the level of fluorescence was normalized to the OD_620nm_ of the same sample. Error bars represent standard deviation of the mean from at least 3 independent biological replicates.

### Graphing and statistics

All statistical analyses and graphing were performed with GraphPad Prism Software version 7.00 for Windows, GraphPad Software, La Jolla California USA, www.graphpad.com.

## Acknowledgments

We thank Richard Johnson, Monica Gupta, Damen Schaak and Kevin Manley for technical expertise and valuable discussions. We are grateful to Guangchun Bai and Marlene Belfort for their generous gifts of antibodies for this work, and to Bill Jacobs and Keith Derbyshire for the gifts of bacterial strains. We also express gratitude to Marlene Belfort and her laboratory members for discussions on intein biology. This work was supported in part by National Institutes of Health grants R21AI104166 and R21AI118351. Additional funding was provided by Health Research Inc grant #11009701.

## References

1. Martin, C.J., Carey, A.F. & Fortune, S.M. A bug’s life in the granuloma. Semin Immunopathol 38, 213–220 (2016).

2. Singh, A. Guardians of the mycobacterial genome: A review on DNA repair systems in Mycobacterium tuberculosis. Microbiology (Reading) 163, 1740–1758 (2017).

3. Boshoff, H.I., Reed, M.B., Barry, C.E., 3rd & Mizrahi, V. DnaE2 polymerase contributes to in vivo survival and the emergence of drug resistance in Mycobacterium tuberculosis. Cell 113, 183–193 (2003).

4. Stephanou, N.C. et al. Mycobacterial nonhomologous end joining mediates mutagenic repair of chromosomal double-strand DNA breaks. J Bacteriol 189, 5237–5246 (2007).

5. Sander, P. et al. Mycobacterium bovis BCG recA deletion mutant shows increased susceptibility to DNA-damaging agents but wild-type survival in a mouse infection model. Infect Immun 69, 3562–3568 (2001).

6. Sassetti, C.M. & Rubin, E.J. Genetic requirements for mycobacterial survival during infection. Proc Natl Acad Sci U S A 100, 12989–12994 (2003).

7. Wipperman, M.F. et al. Mycobacterial Mutagenesis and Drug Resistance Are Controlled by Phosphorylation- and Cardiolipin-Mediated Inhibition of the RecA Coprotease. Mol Cell 72, 152–161 e157 (2018).

8. Gopaul, K.K., Brooks, P.C., Prost, J.F. & Davis, E.O. Characterization of the two Mycobacterium tuberculosis recA promoters. J Bacteriol 185, 6005–6015 (2003).

9. Prasad, D. & Muniyappa, K. The Anionic Phospholipids in the Plasma Membrane Play an Important Role in Regulating the Biochemical Properties and Biological Functions of RecA Proteins. Biochemistry 58, 1295–1310 (2019).

10. Novikova, O., Topilina, N. & Belfort, M. Enigmatic distribution, evolution, and function of inteins. J Biol Chem 289, 14490–14497 (2014).

11. Mills, K.V., Johnson, M.A. & Perler, F.B. Protein splicing: how inteins escape from precursor proteins. J Biol Chem 289, 14498–14505 (2014).

12. Zhang, L., Zheng, Y., Callahan, B., Belfort, M. & Liu, Y. Cisplatin inhibits protein splicing, suggesting inteins as therapeutic targets in mycobacteria. J Biol Chem 286, 1277–1282 (2011).

13. Liu, X.Q. & Yang, J. Prp8 intein in fungal pathogens: target for potential antifungal drugs. FEBS Lett 572, 46–50 (2004).

14. Cheriyan, M. & Perler, F.B. Protein splicing: A versatile tool for drug discovery. Adv Drug Deliv Rev 61, 899–907 (2009).

15. Saves, I., Westrelin, F., Daffe, M. & Masson, J.M. Identification of the first eubacterial endonuclease coded by an intein allele in the pps1 gene of mycobacteria. Nucleic Acids Res 29, 4310–4318 (2001).

16. Davis, E.O., Sedgwick, S.G. & Colston, M.J. Novel structure of the recA locus of Mycobacterium tuberculosis implies processing of the gene product. J Bacteriol 173, 5653–5662 (1991).

17. Pietrokovski, S. Intein spread and extinction in evolution. Trends Genet 17, 465–472 (2001).

18. Nicastri, M.C. et al. Internal disulfide bond acts as a switch for intein activity. Biochemistry 52, 5920–5927 (2013).

19. Topilina, N.I. et al. SufB intein of Mycobacterium tuberculosis as a sensor for oxidative and nitrosative stresses. Proc Natl Acad Sci U S A 112, 10348–10353 (2015).

20. Topilina, N.I., Novikova, O., Stanger, M., Banavali, N.K. & Belfort, M. Post-translational environmental switch of RadA activity by extein-intein interactions in protein splicing. Nucleic Acids Res 43, 6631–6648 (2015).

21. Chong, S., Williams, K.S., Wotkowicz, C. & Xu, M.Q. Modulation of protein splicing of the Saccharomyces cerevisiae vacuolar membrane ATPase intein. J Biol Chem 273, 10567–10577 (1998).

22. Lennon, C.W., Stanger, M., Banavali, N.K. & Belfort, M. Conditional Protein Splicing Switch in Hyperthermophiles through an Intein-Extein Partnership. mBio 9 (2018).

23. Lennon, C.W., Stanger, M. & Belfort, M. Protein splicing of a recombinase intein induced by ssDNA and DNA damage. Genes Dev 30, 2663–2668 (2016).

24. Lennon, C.W., Stanger, M.J. & Belfort, M. Mechanism of Single-Stranded DNA Activation of Recombinase Intein Splicing. Biochemistry 58, 3335–3339 (2019).

25. Davis, E.O., Jenner, P.J., Brooks, P.C., Colston, M.J. & Sedgwick, S.G. Protein splicing in the maturation of M. tuberculosis recA protein: a mechanism for tolerating a novel class of intervening sequence. Cell 71, 201–210 (1992).

26. Martin, D.D., Xu, M.Q. & Evans, T.C., Jr. Characterization of a naturally occurring trans-splicing intein from Synechocystis sp. PCC6803. Biochemistry 40, 1393–1402 (2001).

27. Lennon, C.W., Wahl, D., Goetz, J.R. & Weinberger, J., 2nd Reactive Chlorine Species Reversibly Inhibit DnaB Protein Splicing in Mycobacteria. Microbiol Spectr 9, e0030121 (2021).

28. Woods, D. et al. Conditional DnaB Protein Splicing Is Reversibly Inhibited by Zinc in Mycobacteria. mBio 11 (2020).

29. Kelley, D.S. et al. Mycobacterial DnaB helicase intein as oxidative stress sensor. Nat Commun 9, 4363 (2018).

30. Zhang, L. et al. Binding and inhibition of copper ions to RecA inteins from Mycobacterium tuberculosis. Chemistry 16, 4297–4306 (2010).

31. Davis, E.O., Dullaghan, E.M. & Rand, L. Definition of the mycobacterial SOS box and use to identify LexA-regulated genes in Mycobacterium tuberculosis. J Bacteriol 184, 3287–3295 (2002).

32. Davis, E.O. et al. DNA damage induction of recA in Mycobacterium tuberculosis independently of RecA and LexA. Mol Microbiol 46, 791–800 (2002).

33. Papavinasasundaram, K.G. et al. Slow induction of RecA by DNA damage in Mycobacterium tuberculosis. Microbiology (Reading) 147, 3271–3279 (2001).

34. Movahedzadeh, F., Colston, M.J. & Davis, E.O. Determination of DNA sequences required for regulated Mycobacterium tuberculosis RecA expression in response to DNA-damaging agents suggests that two modes of regulation exist. J Bacteriol 179, 3509–3518 (1997).

35. Ward, S.K., Hoye, E.A. & Talaat, A.M. The global responses of Mycobacterium tuberculosis to physiological levels of copper. J Bacteriol 190, 2939–2946 (2008).

36. Rowland, J.L. & Niederweis, M. A multicopper oxidase is required for copper resistance in Mycobacterium tuberculosis. J Bacteriol 195, 3724–3733 (2013).

37. Bal, W., Sokolowska, M., Kurowska, E. & Faller, P. Binding of transition metal ions to albumin: sites, affinities and rates. Biochim Biophys Acta 1830, 5444–5455 (2013).

38. Eastman, A. Characterization of the adducts produced in DNA by cis-diamminedichloroplatinum(II) and cis-dichloro(ethylenediamine)platinum(II). Biochemistry 22, 3927–3933 (1983).

39. Bruhn, S., L., Toney, Jeffrey, H., Lippard, Stephen, J. Biological Processing of DNA Modified by Platinum Compounds. Progress in Inorganic Chemistry 38, 477–516 (1990).

40. Darwin, K.H., Ehrt, S., Gutierrez-Ramos, J.C., Weich, N. & Nathan, C.F. The proteasome of Mycobacterium tuberculosis is required for resistance to nitric oxide. Science 302, 1963–1966 (2003).

41. Festa, R.A. et al. Prokaryotic ubiquitin-like protein (Pup) proteome of Mycobacterium tuberculosis [corrected]. PLoS One 5, e8589 (2010).

42. Hu, G. et al. Structure of the Mycobacterium tuberculosis proteasome and mechanism of inhibition by a peptidyl boronate. Mol Microbiol 59, 1417–1428 (2006).

43. Muller, A.U., Imkamp, F. & Weber-Ban, E. The Mycobacterial LexA/RecA-Independent DNA Damage Response Is Controlled by PafBC and the Pup-Proteasome System. Cell Rep 23, 3551–3564 (2018).

44. Horii, T., Ozawa, N., Ogawa, T. & Ogawa, H. Inhibitory effects of N- and C-terminal truncated Escherichia coli recA gene products on functions of the wild-type recA gene. J Mol Biol 223, 105–114 (1992).

45. Karlin, S. & Brocchieri, L. Evolutionary conservation of RecA genes in relation to protein structure and function. J Bacteriol 178, 1881–1894 (1996).

46. Guhan, N. & Muniyappa, K. Mycobacterium tuberculosis RecA intein possesses a novel ATP-dependent site-specific double-stranded DNA endonuclease activity. J Biol Chem 277, 16257–16264 (2002).

47. Guhan, N. & Muniyappa, K. The RecA intein of Mycobacterium tuberculosis promotes cleavage of ectopic DNA sites. Implications for the dispersal of inteins in natural populations. J Biol Chem 277, 40352–40361 (2002).

48. Guhan, N. & Muniyappa, K. Mycobacterium tuberculosis RecA intein, a LAGLIDADG homing endonuclease, displays Mn(2+) and DNA-dependent ATPase activity. Nucleic Acids Res 31, 4184–4191 (2003).

49. Wood, D.W., Belfort, M. & Lennon, C.W. Inteins-mechanism of protein splicing, emerging regulatory roles, and applications in protein engineering. Front Microbiol 14, 1305848 (2023).

50. Hill, P.W.S. et al. The vulnerable versatility of Salmonella antibiotic persisters during infection. Cell Host Microbe 29, 1757–1773 e1710 (2021).

51. Salini, S. et al. The Error-Prone Polymerase DnaE2 Mediates the Evolution of Antibiotic Resistance in Persister Mycobacterial Cells. Antimicrob Agents Chemother 66, e0177321 (2022).

52. Tomasz, M. Mitomycin C: small, fast and deadly (but very selective). Chem Biol 2, 575–579 (1995).

53. Drlica, K. & Zhao, X. DNA gyrase, topoisomerase IV, and the 4-quinolones. Microbiol Mol Biol Rev 61, 377–392 (1997).

54. Cumming, B.M. et al. The Physiology and Genetics of Oxidative Stress in Mycobacteria. Microbiol Spectr 2 (2014).

55. Ehrt, S. & Schnappinger, D. Mycobacterial survival strategies in the phagosome: defence against host stresses. Cell Microbiol 11, 1170–1178 (2009).

56. Newton, G.L., Buchmeier, N. & Fahey, R.C. Biosynthesis and functions of mycothiol, the unique protective thiol of Actinobacteria. Microbiol Mol Biol Rev 72, 471–494 (2008).

57. Stafford, S.L. et al. Metal ions in macrophage antimicrobial pathways: emerging roles for zinc and copper. Biosci Rep 33 (2013).

58. Lee, A.K., Detweiler, C.S. & Falkow, S. OmpR regulates the two-component system SsrA-ssrB in Salmonella pathogenicity island 2. J Bacteriol 182, 771–781 (2000).

59. Groisman, E.A. The pleiotropic two-component regulatory system PhoP-PhoQ. J Bacteriol 183, 1835–1842 (2001).

60. Schuhmacher, D.A. & Klose, K.E. Environmental signals modulate ToxT-dependent virulence factor expression in Vibrio cholerae. J Bacteriol 181, 1508–1514 (1999).

61. Hase, C.C. & Mekalanos, J.J. TcpP protein is a positive regulator of virulence gene expression in Vibrio cholerae. Proc Natl Acad Sci U S A 95, 730–734 (1998).

62. DiRita, V.J. Co-ordinate expression of virulence genes by ToxR in Vibrio cholerae. Mol Microbiol 6, 451–458 (1992).

63. Larsen, M.H. et al. Efficacy and safety of live attenuated persistent and rapidly cleared Mycobacterium tuberculosis vaccine candidates in non-human primates. Vaccine 27, 4709–4717 (2009).

64. Sambandamurthy, V.K. et al. Long-term protection against tuberculosis following vaccination with a severely attenuated double lysine and pantothenate auxotroph of Mycobacterium tuberculosis. Infect Immun 73, 1196–1203 (2005).

65. Jain, P. et al. Specialized transduction designed for precise high-throughput unmarked deletions in Mycobacterium tuberculosis. mBio 5, e01245–01214 (2014).

66. Cobbert, J.D. et al. Caught in action: selecting peptide aptamers against intrinsically disordered proteins in live cells. Sci Rep 5, 9402 (2015).

67. Schmittgen, T.D. & Livak, K.J. Analyzing real-time PCR data by the comparative C(T) method. Nat Protoc 3, 1101–1108 (2008).

68. Roy, A., Kucukural, A. & Zhang, Y. I-TASSER: a unified platform for automated protein structure and function prediction. Nat Protoc 5, 725–738 (2010).

69. Yang, J. et al. The I-TASSER Suite: protein structure and function prediction. Nat Methods 12, 7–8 (2015).

70. Zhang, Y. I-TASSER server for protein 3D structure prediction. BMC Bioinformatics 9, 40 (2008).

71. Datta, S., Ganesh, N., Chandra, N.R., Muniyappa, K. & Vijayan, M. Structural studies on MtRecA-nucleotide complexes: insights into DNA and nucleotide binding and the structural signature of NTP recognition. Proteins 50, 474–485 (2003).

