## Supplemental info for "Conditional protein splicing of the *Mycobacterium tuberculosis* RecA intein in its native host"

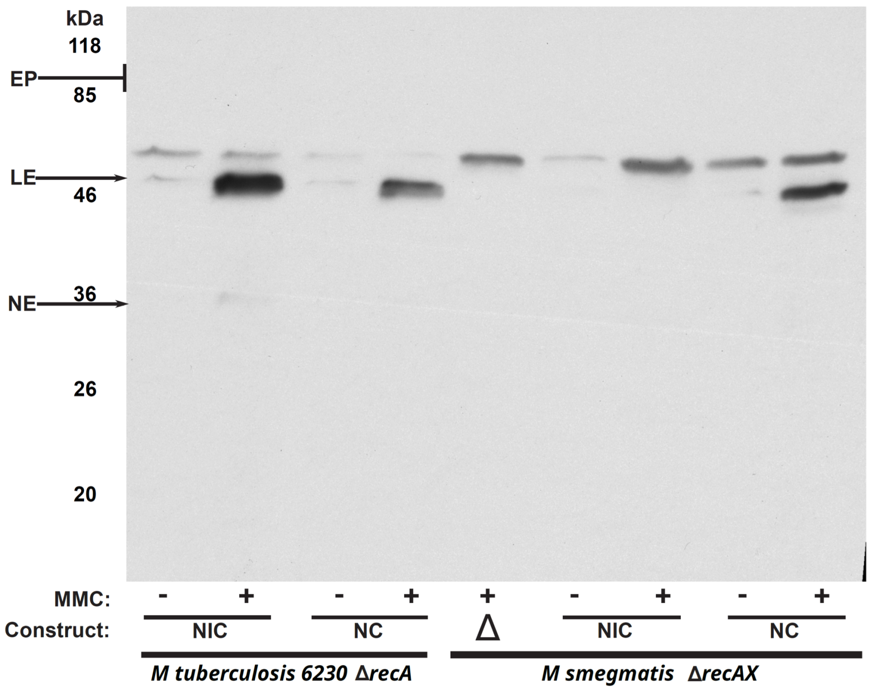


**Supplemental Figure S1:** RecA precursor was not detectable in other mycobacteria. *M. tuberculosis ΔrecA* and *M. smegmatis ΔrecAX* were complemented with intein-containing *recA* (NIC) or inteinless *recA* (NC) alleles driven by their native mycobacterial promoters. Cells were grown to mid-log and treated for 3 days with Mitomycin C to induce *recA* transcription and RecA production. Protein from cultures was harvested for western blotting against the N-extein, which detects ligated exteins (LE) and the N-extein (NE) as well as precursor (see Figure 1B). EP: expected size of precursor RecA, which was not detected.

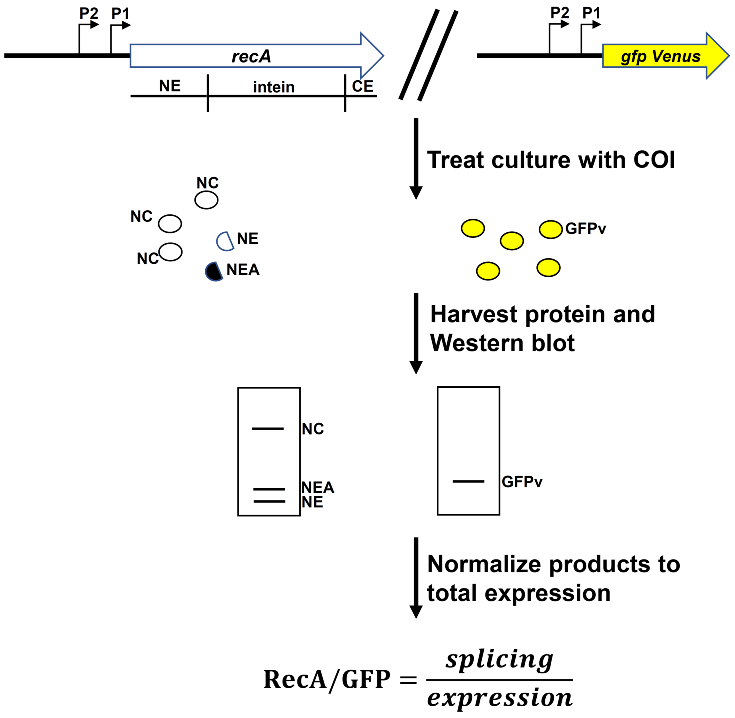


**Supplemental Figure S2:** Schematic of experimental setup and analysis. Mtb auxotroph mc$^2$6230 harboring a native *recA* allele and a *gfp* allele driven by the native Mtb *recA* promoters is grown to the desired phase and treated with conditions of interest (COI). If treatment induces *recA* production, RecA products and GFP will be made at a higher level compared to untreated. After treatment, total protein lysate is extracted and subjected to western blotting targeting the N-extein (left), GFP (right) or Intein (not shown). Western blots are quantitated by densitometry and the ratio of RecA product to GFP is calculated to normalize for changes in transcriptional expression.

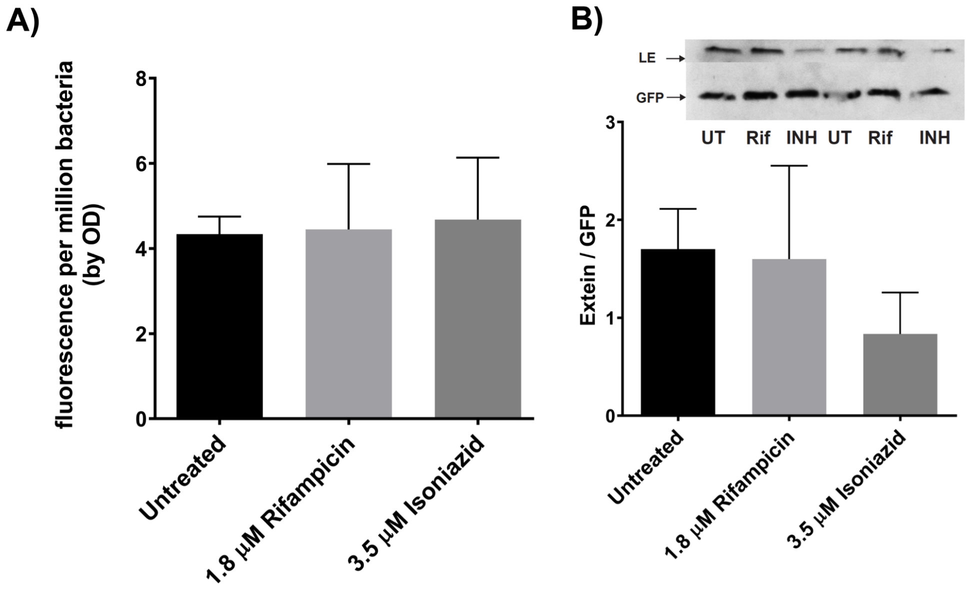


**Supplemental Figure S3:** First-line therapeutics rifampicin and Isoniazid do not induce the *recA* transcription or RecA splicing. Mtb auxotrophic strain mc$^2$6230 harboring our *recA* transcriptional reporter was grown to mid-log phase and then treated with rifampicin or isoniazid. (A) *recA* transcription in response to the two therapeutics as read by our GFP based transcriptional reporter system. (B) Western blot of RecA production in response to rifampicin or isoniazid. Error bars represent standard deviation of three biological repeats.

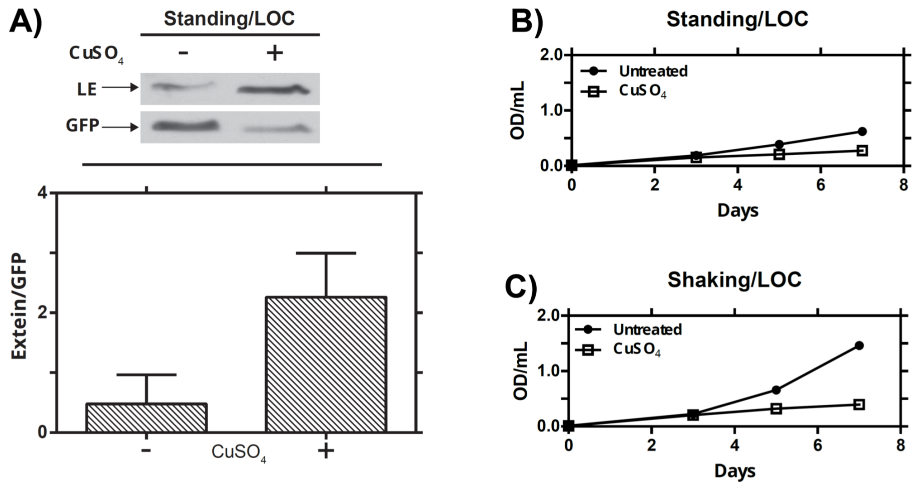


**Supplemental Figure S4:** Growth does not affect copper-induced splicing. Mtb was grown in the presence or absence of copper for seven days either standing or shaking in low oxygen conditions supplemented with carbon dioxide (LOC). A) Representative western blot with quantitative analysis of RecA ligated extein (LE) produced in response to copper in standing LOC. Error bars represent standard deviation of three biological repeats. B) Optical density of cultures grown in standing LOC conditions with copper absent (black circle) or present (open square). C) Optical densities of cultures grown in shaking LOC conditions with copper absent (black circle) or present (open square).

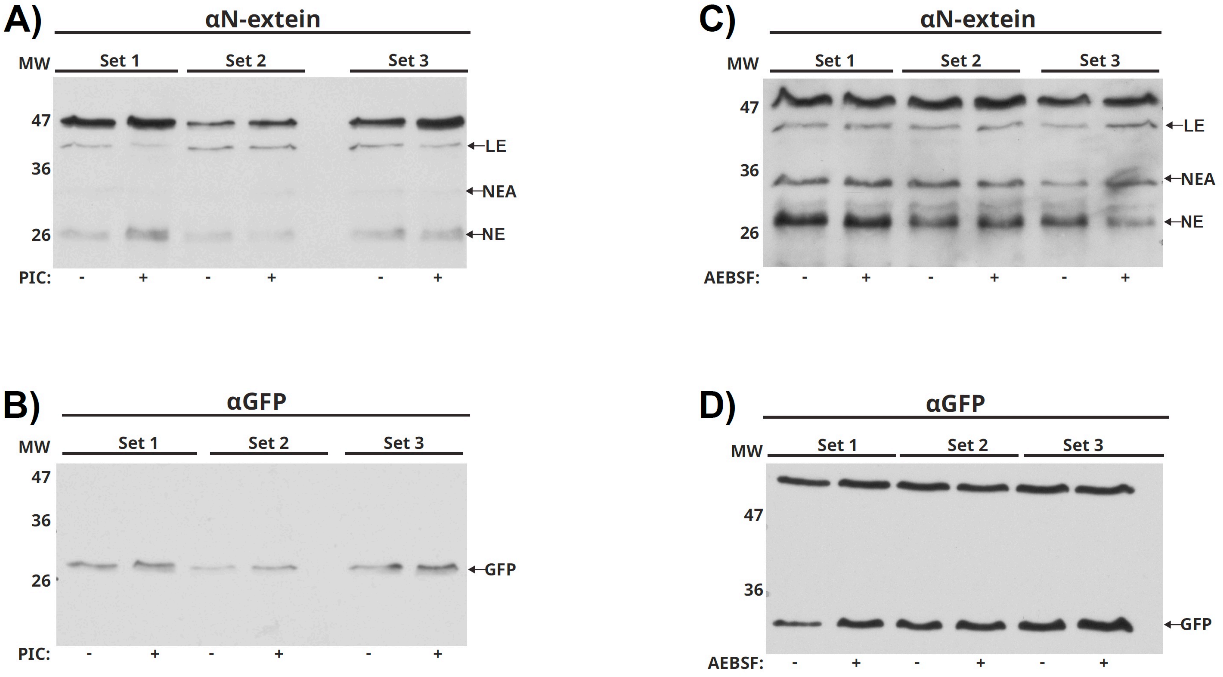


**Supplemental Figure S5:** Treatment with protease inhibitors does not affect levels of RecA or intein. Mtb mc[2]6230 harboring a native *recA* allele and a *gfp* allele driven by the native Mtb *recA* promoters was grown to (A, B) mid-log phase before treating with 1:1000 dilution of Sigma Aldrich's protease inhibitor cocktail for 24hrs in triplicate. (A) Western blot analysis targeting the N-extein. (B) Western blot analysis targeting GFP. (C-D) Mtb mc[2]6230 harboring a native *recA* allele and a GFP allele driven by the native *recA* promoters was grown to late-log phase and treated with AEBSF for 48hrs in triplicate. Western blot analysis targeted the N-extein (C) or GFP (D).

**Supplemental Table S1: Strains and plasmids used in this study.**

| Strain | Description | Source |
|---|---|---|
| $mc^2$6230 | Mtb H37Rv (*ΔpanCD*, *ΔRD1*) | Gift from Dr. Bill Jacobs [1] |
| $mc^2$6230*ΔrecA* | *recA* knockout (*ΔrecA*, *ΔpanCD*, *ΔRD1*) | This Paper |
| *Mycobacterium smegmatis* $mc^2$155 | WT | Acquired from BEI [2] |
| *Mycobacterium smegmatis ΔrecAX* | $mc^2$155 *recAX* knockout (*ΔrecAX*) | Gift from Dr. Keith Derbyshire |
| *E. coli BLR* | *ΔrecA* derivative of BL21 | Novagen |
| *E. coli BL21* | For protein expression and purification | Novagen |

**Supplemental Table S2: Plasmids used in this study.**

| Designation | Description | Source |
|---|---|---|
| pMBC409 | Single Copy, integrating (AttP) vector to express transcriptional reporters or *recA* complements; $Kan^R$, $Hyg^R$, $Amp^R$ | Described in Girardin et al [3] |
| pMBC1809 | mVenus-based *recA* transcriptional reporter; $Kan^R$, $Hyg^R$, $Amp^R$ | This Paper |
| pMBC2149 | Full Mtb *recA* gene driven by native Mtb promoters; $Kan^R$, $Hyg^R$, $Amp^R$ | This Paper |
| pMBC2169 | Inteinless Mtb *recA* gene driven by native Mtb promoters; $Kan^R$, $Hyg^R$, $Amp^R$ | This Paper |
| pMBC2290 | Mtb N-extein driven by native Mtb promoters; $Kan^R$, $Hyg^R$, $Amp^R$ | This Paper |
| pMBC1650 | pET28a+ expressing N-extein; $Kan^R$ | This Paper |
| pMBC1651 | pET28A+ expressing intein; $Kan^R$ | This Paper |
| pMBC1652 | pET28a+ expressing C-extein; $Kan^R$ | This paper |

**Supplemental Table S3: Primers used in this study.**

| Designation | Sequence | Description |
|---|---|---|
| KM4059 | gggggatcctctagatttaagaaggagatatacatatggtgagcaagggcgaggagctg | mVenus forward |
| KM4138 | gggaagctttgatcaccgcggccatg | mVenus reverse |
| KM4194 | tgcagtggatcccgcaccgccgccagg | *recA* promoter forward |
| KM4195 | cctagtggatcccatggtgcctctcctgtg | *recA* promoter reverse |
| KM4345 | agagatatacggcccggagt | *recA* qRT |
| KM4346 | ccgagcttcttggcatagtc | *recA* qRT |
| KM4290 | acgtaaacggccacaagttc | GFPv qRT |
| KM4291 | aagtcgtgctgcttcatgtg | GFPv qRT |
| KM3727 | gggaattcacgcagacccccgatcgg | *recA* NE for protein purification fwd |
| KM3728 | ggaagctttcacttgttcttgacgaccttgac | *recA* NE for protein purification rev |
| KM3729 | gggaattctgcctcgcagagggc | *recA* intein for protein purification fwd |
| KM3730 | ggaagctttcaacagttgtgcacgacaaccc | *recA* intein for protein purification rev |
| KM3731 | gggaattctcgcccccccttcaagca | *recA* CE for protein purification fwd |
| KM3732 | gggagctctcagaagtcgacggggg | *recA* CE for protein purification rev |
| KM5105 | gatatcgtgttgagcagatcgtcggtgatccgga | *recA* full complement fwd |
| KM5106 | gcggccgctcagaagtcgacgggggcgg | *recA* full complement rev |
| KM5584 | gatatcgcctgctcttcgcgctcagaag | *recA* inteinless complement NE fwd |
| KM5585 | caaggtcgtcaagaacaagtgttcgccccccttcaagcagg | *recA* inteinless complement NE rev |
| KM5586 | gaagggggggcgaacacttgttcttgacgaccttgacccggg | *recA* inteinless complement CE fwd |
| KM5587 | gatatcgacgccgaaaggtcagatccgg | *recA* inteinless complement CE rev |
| KM6016 | aagcttgagcagatcgtcggtgatc | *recA* N-Extein only complement fwd |
| KM6017 | aagcttctacttgttcttgacgaccttgac | *recA* N-Extein only complement rev |

**Supplemental References:**


1. Sambandamurthy, V.K. *et al. Mycobacterium tuberculosis* DeltaRD1 DeltapanCD: a safe and limited replicating mutant strain that protects immunocompetent and immunocompromised mice against experimental tuberculosis. *Vaccine* **24**, 6309-6320 (2006).
2. Snapper, S.B., Melton, R.E., Mustafa, S., Kieser, T. & Jacobs, W.R., Jr. Isolation and characterization of efficient plasmid transformation mutants of *Mycobacterium smegmatis*. *Mol Microbiol* **4**, 1911-1919 (1990).
3. Girardin, R.C. & McDonough, K.A. Small RNA Mcr11 requires the transcription factor AbmR for stable expression and regulates genes involved in the central metabolism of *Mycobacterium tuberculosis*. *Mol Microbiol* **113**, 504-520 (2020).
